# *Streptococcus pyogenes* Cas9 ribonucleoprotein delivery for efficient, rapid and marker-free gene editing in *Trypanosoma* and *Leishmania*

**DOI:** 10.1101/2023.10.25.563974

**Authors:** Asencio Corinne, Hervé Perrine, Morand Pauline, Oliveres Quentin, Morel Chloé Alexandra, Prouzet-Mauleon Valérie, Biran Marc, Monic Sarah, Bonhivers Mélanie, Robinson Derrick Roy, Ouellette Marc, Rivière Loïc, Bringaud Frédéric, Tetaud Emmanuel

## Abstract

Kinetoplastids are unicellular eukaryotic flagellated parasites found in a wide range of hosts within the animal and plant kingdoms. They are known to be responsible in humans for African sleeping sickness (*Trypanosoma brucei*), Chagas disease (*Trypanosoma cruzi*), and various forms of leishmaniasis (*Leishmania* spp.), as well as several animal diseases with important economic impact (African trypanosomes, including *T. congolense*). Understanding the biology of these parasites necessarily implies the ability to manipulate their genomes. In this study, we demonstrate that transfection of a ribonucleoprotein complex, composed of recombinant *Streptococcus pyogenes* Cas9 (*Sp*Cas9) and an *in vitro*-synthesized guide RNA, results in rapid and efficient genetic modifications of trypanosomatids, in marker-free conditions. This approach was successfully developed to inactivate, delete and mutate candidate genes in various stages of the life cycle of *T. brucei* and *T. congolense*, and *Leishmania* promastigotes. The functionality of *Sp*Cas9 in these parasites now provides, to the research community working on these parasites, a rapid and efficient method of genome editing, without requiring plasmid construction and selection by antibiotics. Importantly, this approach is adaptable to any wild-type parasite, including field isolates.

## INTRODUCTION

In the realm of modern molecular biology, few innovations have captured the world’s attention like CRISPR-Cas9. The CRISPR-Cas9 system has catapulted molecular biology into a new era of precision gene editing and genetic engineering. CRISPR (Clustered Regularly Interspaced Short Palindromic Repeats) and CRISPR-associated protein 9 (CRISPR-Cas9) is a revolutionary genome-editing technology that has unlocked unprecedented opportunities for precise manipulation of the genetic code. Since its discovery in the early 2000s (Mojica et al., 2005, Barrangou et al., 2007), CRISPR-Cas9 has rapidly evolved into a game-changing tool with the potential to transform medicine, agriculture, biotechnology, and various other fields (Jinek et al., 2012). This groundbreaking system is derived from a remarkable natural defense mechanism found in bacteria and archaea, allowing researchers to edit, correct, or modify genes with an accuracy and efficiency that was previously unimaginable (Ishino et al., 1987, Mojica et al., 2005, Barrangou et al., 2007).

Obviously, this technology was very quickly used to study trypanosomatids, which are unicellular eukaryotic flagellate parasites that affect millions of people and animals worldwide. In humans, they are responsible for African sleeping sickness (Human African trypanosomiasis), Chagas disease (American trypanosomiasis), and leishmaniasis in South America, Africa, India, the Mediterranean region and the Middle East. African Animal Trypanosomiasis, including Nagana, are debilitating diseases affecting livestock, primarily cattle in sub-Saharan Africa. Nagana, which is caused by *Trypanosoma congolense*, *Trypanosoma vivax* and *Trypanosoma burcei brucei*, leads to severe health issues, including anemia, weight loss, and decreased productivity in infected animals, making it a significant economic and agricultural concern in affected regions (Desquesnes et al., 2022). Understanding the biology of these parasites necessarily requires the ability to manipulate their genomes. Before the development of the CRISPR-Cas9 system, genome modification was achieved through homologous recombination with resistance markers for selection, or by RNA interference, but only in *T. brucei* (Ngo et al., 1998) and *T. congolense* (Inoue et al., 2002, Coustou et al., 2012) as *T. cruzi* and *Leishmania* spp do not share the RNA interference machinery (Kolev et al., 2011). Because of genome diploidy, deletion/inactivation of genes proved to be time-consuming and conditioned by the limited number of available antibiotic resistance markers.

The CRISPR-Cas9 system has been now employed to modify the genomes of *T. brucei*, *T. cruzi* and *Leishmania* spp. with a rapidly growing number of publications (founding articles are : (Peng et al., 2014, Sollelis et al., 2015, Zhang and Matlashewski, 2015, Zhang et al., 2017, Lander et al., 2015, Lander et al., 2016, Lander et al., 2017, Beneke et al., 2017, Rico et al., 2018, Shaw et al., 2020, Kovarova et al., 2022). In most studies, Cas9 or Cas9-gRNA complexes are endogenously expressed after transfection of the parasites and selection with antibiotic resistance markers. However, the CRISPR-Cas9 system still has some shortcomings, such as the impact of constitutive Cas9 expression, which result in genome instability (Zhang et al., 2017, Boutin et al., 2021) and can lead to a decrease in cell growth (Ryan et al., 2014, Peng et al., 2014). Conditional expression of Cas9, as described in *T. brucei*, is helpful to overcome this issue (Rico et al., 2018, Kovarova et al., 2022). Another issue with the CRISPR-Cas9 system is the potential off-target genome disruption (Fu et al., 2013). The ongoing development of new, increasingly accurate Cas9 variants, should also limit this problem. Interestingly, CRISPR-mediated editing can be achieved by transfecting cells with *in vitro*-generated Cas9 protein/guide RNA complexes. As recombinant Cas9 is rapidly eliminated after transfection, this approach, which limits potential off-target (D’Astolfo et al., 2015) and reduces toxicity issues (Kim et al., 2014, Liang et al., 2015), has proven to be particularly efficient in *T. cruzi* and *Plasmodium* (Soares Medeiros et al., 2017, Crawford et al., 2017). This method is straightforward, fast, and highly efficient, as it does not require gene cloning and minimizes the use of selection markers. However, in *T. cruzi*, *Leishmania* spp. and *T. brucei*, this approach relies exclusively on the use of a smaller Cas9, isolated from *Staphylococcus aureus* (*Sa*Cas9) (Soares Medeiros et al., 2017). The same authors also reported that Cas9 isolated from *Streptococcus pyogenes* (*Sp*Cas9), despite being the most commonly used and commercialized, does not function in these parasites, and the lack of activity appears to be due to the larger size of *Sp*Cas9 compared to *Sa*Cas9, which is approximately 40 kDa smaller (Soares Medeiros et al., 2017).

In this study, we have demonstrated that *Sp*Cas9 is fully functional after transfection of the Cas9/gRNA complex in both the bloodstream (BSF) and procyclic (PCF) forms of *T. brucei* and *T. congolense*, as well as in the promastigotes of *Leishmania infantum*. This Cas9/gRNA complex can be delivered into cells, with or without a repair sequence, to inactivate, mutate, or tag candidate genes, without the need for selection markers and with very high efficiency. The functionality of *Sp*Cas9 in these parasites now provides the research community working on these parasites a rapid and efficient method for genome edition without requiring gene cloning and/or selection, that only requires cloning of modified cell lines. It will also allow genome editing of cells that are difficult to cultivate or whose cell density is too low to envisage using conventional techniques, such as field isolates. Finally, this approach should be adaptable to all kinetoplastids and, importantly, any transfectable cell type.

## RESULTS

### The size of the *Sp*Cas9/RNP complex has no detectible impact on genome editing activity

Soares Medeiros *et al*. described in *T. cruzi*, *T. brucei* and *Leishmania* the unexpected result that exogenous ribonucleoprotein Cas9 from *Streptococcus pyogenes* (*Sp*Cas9, 163 kDa) was not functional after transfection, in contrast to the smaller *Staphylococcus aureus* Cas9 (*Sa*Cas9, 124 kDa) (Soares Medeiros et al., 2017). Since *Sp*Cas9 is fully active when endogenously expressed (Peng et al., 2014), it was proposed that ribonucleoprotein *Sp*Cas9 complexes are not internalized by electroporation in trypanosomatids. To revisit these data, we tried to inactivate a constitutively expressed cytosolic GFP in *T. brucei* PCF using extracellular RNP complexes composed of *Sp*Cas9 and guide RNAs (gRNAs). We used the commercial *Sp*Cas9 from Integrated DNA Technologies (IDT, *Sp*Cas9 nuclease V3) and three gRNAs targeting different sequences of the GFP gene (**Table 1**). GFP expression was monitored by flow cytometry at 24, 48, 72, and 144 h post-transfection with the RNP complexes (**Figure 1A**). 5×10^5^ GFP-expressing *T. brucei* PCF cells were subjected to electroporation with 20 µg of gRNA-loaded *Sp*Cas9 (GFP1, 2 or 3), or gRNA only as a control (GFP2). Unexpectedly, transfection of GFP-expressing PCF without *Sp*Cas9 (**Figure 1A - Top left panel**) resulted in a very strong drop in fluorescence after 24 h for about 50% of the population, probably related to the electroporation conditions. However, 48 h later, GFP expression recovered to its initial level (**Figure 1A - Top left panel**). Importantly, a strong decrease in fluorescence occurred immediately after transfection with GFP1, GFP2 and GFP3 gRNA-loaded *Sp*Cas9, which did not recover to its initial level after 72 h (**Figure 1A – gRNA GFP1, GFP2 and GFP3**), indicating that a significant portion of cells do not express active GFP anymore. It is noteworthy that GFP1 and GFP2 gRNA are more efficient at inactivating GFP expression than GFP3 gRNA (**Figure 1A – Bar chart**) and the efficiency depends on the amount of RNP complex transfected (1.5-fold increase of GFP-negative cells with a 3-fold increase of GFP2 gRNA-loaded *Sp*Cas9) (**Figure 1A – Bar chart**).

**Figure 1-.**
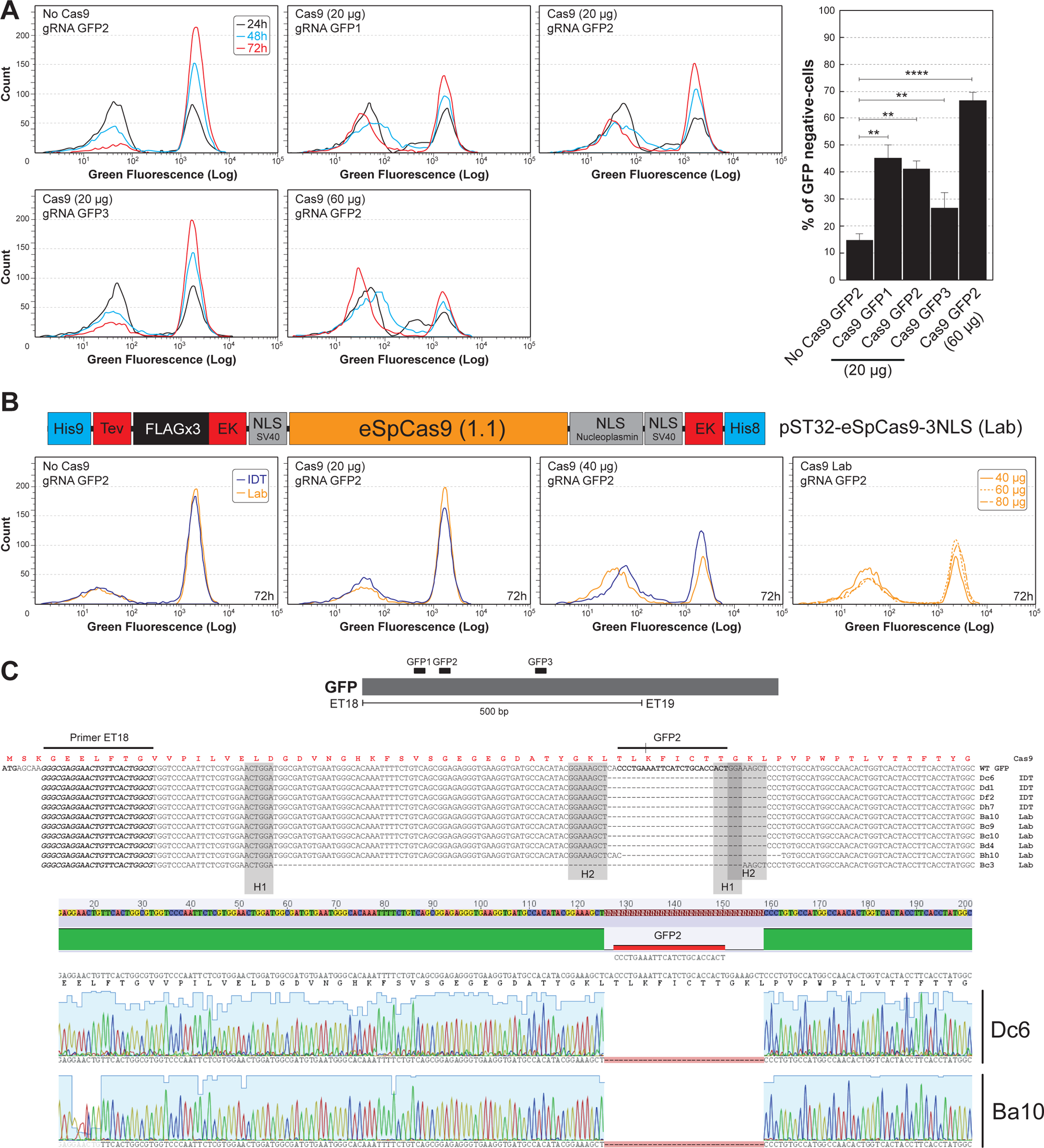
GFP inactivation in *T. brucei* PCF. **(A)** Fluorescence flow cytometry analysis of *T. brucei* constitutively expressing a cytosolic GFP. GFP fluorescence was monitored over time from 24 to 72 h after transfection with 20 µg (no Cas9, Cas9/gRNA GFP1, Cas9/gRNA GFP2, Cas9/gRNA GFP3) or 60 µg (Cas9/gRNA GFP2) of RNP complexes from IDT, and a bar chart showing the percentage of GFP-negative cells at 72 to 144 h after transfection with the different guides. **(B)** The top panel shows a schematic representation of the plasmid allowing e*Sp*Cas9 expression in *E. coli*. The blue boxes represent the two polyhistidine sequences at the N and C-termini of the protein, the red boxes represent the cleavage sites of TEV and enterokinase (EK) proteases, the gray boxes represent the 3 nuclear localization signals (NLS), the black box represents 3 repeats of the FLAG epitope and the orange box represents the e*Sp*Cas9 coding sequence. The bottom panel shows the fluorescence flow cytometry analysis of *T. brucei* expressing the GFP monitored at 72 h after transfection with RNPs complexes from IDT or laboratory-purified (Lab) (no Cas9, 20 µg Cas9/gRNA GFP2, 40 µg Cas9/gRNA GFP2, 40, 60 and 80 µg Cas9/gRNA GFP2). **(C)** Sequence comparison of a portion of the GFP gene from clones no longer expressing GFP. The sequence shows only the region targeted by the GFP2 guide RNA. The gray boxes (H1 and H2) highlight the homology regions probably used for repair by MMEJ. Sequences resulting from inactivation by laboratory-purified Cas9 and those from commercial Cas9 are labeled Lab and IDT respectively. Below is shown the corresponding chromatogram of the Dc6 and Ba10 clones.

**Table 1-.** Summary of the various CRISPR/Cas9 inactivation experiments.

| Gene targeted | Parasite | Inactivation cassette | Numbers of tested clones | Homozygous | Heterozygous | Both alleles inactivated (%) |
| --- | --- | --- | --- | --- | --- | --- |
| **Fis1** | *T. brucei* PCF | *BleR* + 3'/5' UTR | 5 | 5 | 0 | 100 |
|  | *T. brucei* BSF |  | 12 | 1 | 11 | 8 |
|  | *T. brucei* BSF | *PacR* without UTR | 34 | 0 | 33 | 0 |
|  | *T. brucei* BSF | *SBS* | 76 | 2 | 3 | 3 |
| **GK** | *T. brucei* PCF | *BleR* + 3'/5' UTR | 11 | 11 | 0 | 100 |
| **ALDH** | *L. infantum* Pro | *PacR* without UTR | 39 | 12 | 1 | 31 |
|  | *L. infantum* Pro | *mRED* without UTR | 6 | 5 | 1 | 83 |
|  | *L. infantum* Pro | *SBS* | 12 | 4 | 0 | 33 |
| **LysoPLA** | *T. congolense* BSF | *BleR* + 3'/5' UTR | 4 | 4 | 0 | 100 |
|  | *T. congolense* PCF | *SBS* | 5 | 3 | 0 | 60 |
| **GK** | *T. congolense* BSF | *BleR* + 3'/5' UTR | 4 | 3 | 1 | 75 |

*SBS*: short sequence containing a succession of stop codons

To confirm that the decrease in GFP expression is indeed caused by Cas9/gRNA-dependent GFP gene inactivation, we cell-sorted and cloned cells failing to express GFP and sequenced the DNA region targeted by the GFP2 guide RNA used (**Figure 1C**). As expected, a 33-bp deletion at the gRNA targeting site was observed in all GFP-negative cells (**Figure 1C**). The deleted region is flanked by two 8-bp homologous sequences, which suggests a repair of double-strand breaks by the microhomology-mediated end-joining (MMEJ) pathway as previously described in *T. cruzi* and *Leishmania* (Peng et al., 2014, Zhang et al., 2017). These data show that the commercial *Sp*Cas9 from IDT is fully functional in our experimental protocols and suggest that the size of the exogenous ribonucleoprotein complex is not a limiting factor, contrary to the hypothesis made for *T. cruzi*. The only difference that we could identify between the IDT *Sp*Cas9 and the one used by Soares Medeiros *et al*. is the number of nuclear localization signals (NLS), three *versus* two, suggesting that the lack of activity in *T. cruzi* could be explained by weak nuclear targeting of the complex after transfection.

In order to produce our own in-house recombinant Cas9 protein, we constructed a recombinant DNA *SpC*as9 sequence (e*Sp*Cas9, a rationally engineered Cas9 with improved specificity (Slaymaker et al., 2016)) containing three NLS regions, *i.e.*, one at the N-terminus and two at the C-terminus of the protein, plus two polyhistidine tracts to allow the purification of the expressed recombinant protein by chromatography (**Figure 1B**). The protein was expressed and purified in *E. coli* (Cf. Experimental procedures, **Figure S1**) and its activity was assayed by replacing the IDT *Sp*Cas9 with the e*Sp*Cas9 in the GFP2 gRNA/Cas9 complex. As expected, the purified e*Sp*Cas9 is able to dose-dependently inactivate GFP expression (**Figure 1B**). Note that above 40 µg of e*Sp*Cas9, there is no further decrease in GFP expression (**Figure 1B, right**). Sequencing of the GFP gene in cloned cell lines confirmed the inactivation of the gene by a 33-bp deletion. A larger 97-bp deletion flanked by two 6-bp homologous regions was also detected in clone Bc3 (**Figure 1C**). It is worth noting that the presence of polyhistidine tracts, one each at the N and C-terminal of the protein, does not appear to affect Cas9 activity. We concluded that the edition of PCF *T. brucei* genes with *Sp*Cas9/gRNA complexes is functional and the use of laboratory-produced e*Sp*Cas9 is equally effective as the commercial *Sp*Cas9 (IDT).

### Development of a marker-free approach for editing the genomes of *T. brucei* PCF and BSF

Cas9-mediated double-strand DNA breaks can be repaired by homology-directed repair (HDR), as long as an appropriate repair template is provided (Peng et al., 2014). We have tested this by inactivating the *TbFis1* gene (Tb927.10.8660), in both *T. brucei* PCF and BSF using marker-dependent and marker-free approaches. *Tb*Fis1 protein is a potential homologue of the mitochondrial fission factor identified in yeast, Fis1p (Mozdy et al., 2000), which enables the recruitment of the dynamin Dnm1 to the mitochondria and triggers mitochondrial fission. The first approach consists of inserting a repair template encoding a resistance gene (in this case against phleomycin, *Ble*R) flanked by 5’ and 3’ regulatory sequences and a short homology region of the *TbFis1* gene flanking the Cas9-cleavage site (50 bp) (**Figure 2A**). Cells were cloned after 8 to 12 days of culture in the presence of phleomycin. The correct insertion of the repair DNA fragment was controlled by PCR and sequencing, as shown in **Figure 2B/C**. After Cas9-mediated recombination, the size of the targeted *TbFis1* gene increased by 880 bp, which corresponds to the size of the repair cassette (**Figure 2B**). Both alleles encoding *Tb*Fis1 (homozygotes) were targeted in 100% of PCF clones but in only 10% of BSF clones (**Figure 2B**). The 90% remaining BSF clones had a single allele inactivated (heterozygotes) (**Table 1**).

**Figure 2-.**
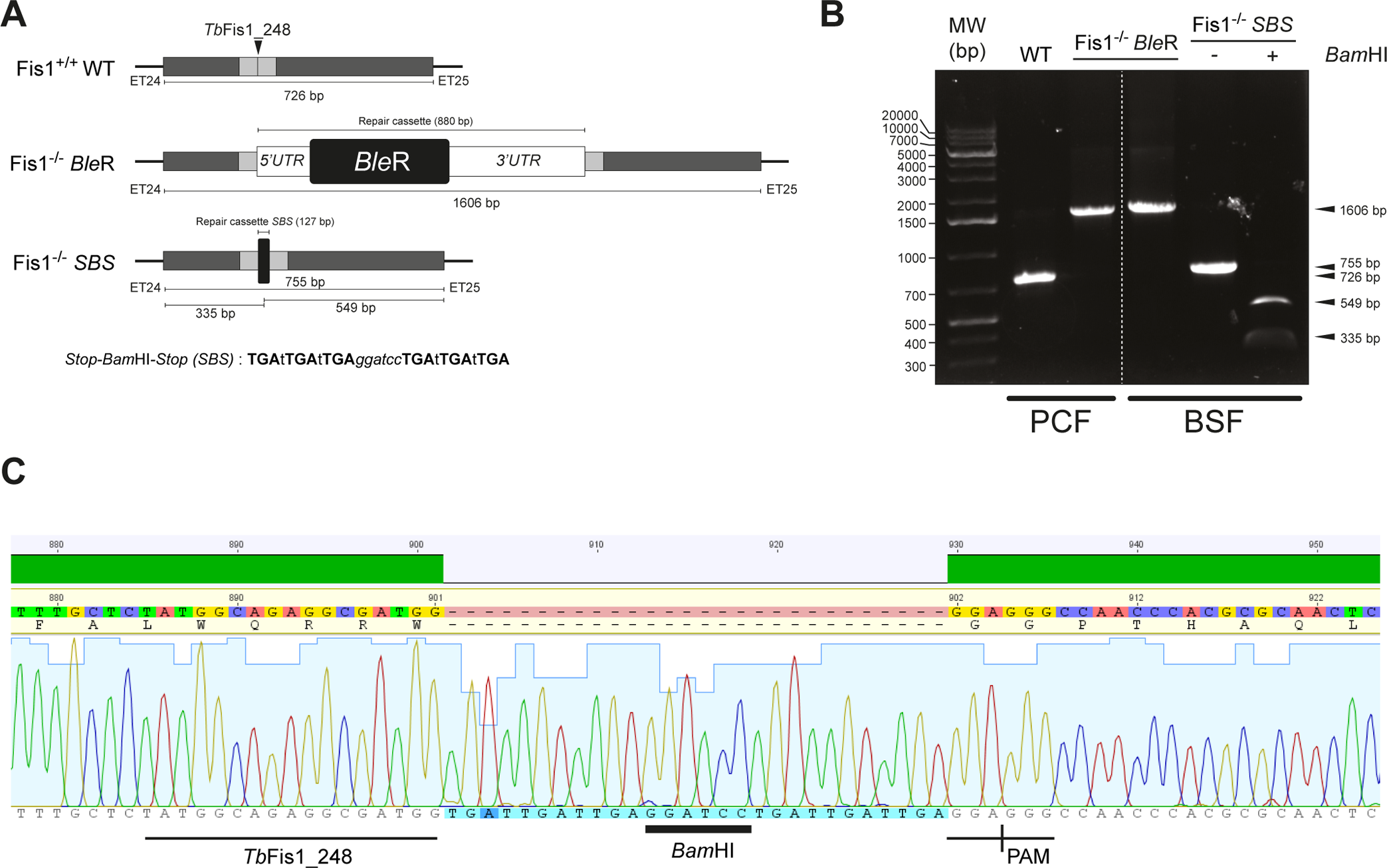
Inactivation of the *TbFis1* gene in both *T. brucei* PCF and BSF. **(A)** Schematic representation of the *Tb*Fis1 locus and the two inactivation strategies by inserting the phleomycin resistance marker (*Ble*R) or a short sequence containing a succession of stop codons (SBS). The position of the guide RNA is indicated by a vertical arrow (*Tb*Fis1_248) and the 50-bp flanking sequences allowing repair through HDR are shown in gray. **(B)** PCR confirmation of *TbFis1* gene inactivation on both alleles in PCF and BSF cells. PCR products (primer ET24/ET25) are directly analyzed on agarose gel (Fis1^−/−^ *Ble*R) or after its digestion with *Bam*HI (Fis1^−/−^ SBS), allowing easy discrimination of gene inactivation on both alleles. **(C)** Chromatogram of the *Tb*Fis1 sequence highlighting the insertion of the SBS cassette.

The second approach consists of inserting a shorter repair cassette composed of stop codons in all three reading frames and a restriction site absent in the targeted gene (here *Bam*HI), flanked by short regions of homology corresponding to 50-bp flanking the Cas9-cleavage site (**Figure 2A - SBS**). The *Bam*HI restriction site enables rapid discrimination of repair template integration on one or two alleles after PCR and digestion with *Bam*HI. Under these marker-free conditions, cells were cloned one to three days after transfection and insertion of the repair cassette was tested by PCR/*Bam*HI digestion and sequencing (**Figure 2B – SBS**). 3% and 4% of the BSF clones tested were inactivated on both alleles or one allele only, respectively (**Table 1**). Obtaining a large number of homozygous mutants confirms that the *TbFis1* gene is not essential for PCF and BSF growth. Incidentally, the inactivation of the *TbFis1* gene did not induce a change in mitochondrial structure (**Figure S2**).

### Inactivation of a *T. brucei* multigene family by *Sp*Cas9

Genetic manipulation of trypanosomatids is often made difficult when high number of resistance markers are required, as exemplified for sequential inactivation of several genes or when multigene families are addressed (Reis-Cunha et al., 2018). We therefore tested whether transfection, with the gRNA/RNP complex, is also effective for the inactivation of a multigene family, here the one encoding glycerol kinase (GK) in *T. brucei* PCF. RNAi-mediated down-regulation of GK expression has shown that, in standard growth conditions, this non-essential gene family is required to metabolize glycerol in *T. brucei* PCF and BSF (Pineda et al., 2018, Allmann et al., 2021). GK is encoded by eleven tandemly-arranged copies distributed over the two alleles, containing five and six copies, respectively (**Figure S2**). We tested a single gRNA targeting the entire multigenic family and a repair cassette including the phleomycin resistance marker (*Ble*R) (**Figure 3A**). The fate of the allelic GK gene clusters after transfection and selection with phleomycin was tested by PCR using primers flanking the Cas9 recognition site. All the tested phleomycin-resistant clones (eleven clones) showed the presence of a single ∼1,500-bp band corresponding to the parental allele (616 bp) inactivated by insertion of the repair cassette (880 bp), suggesting that GK genes are inactivated (**Figure 3B**, **Table 1**). This was confirmed by a Southern-blot analysis with the GK probe after digestion of the genomic DNA with a restriction enzyme present once in each of the GK repeat unit (*Kpn*I), which generated a 5,197-bp band corresponding to the GK copy located at the 5’ extremity of both allelic clusters (two copies) plus an intense 3,582-bp band corresponding to the other GK genes (nine copies) (**Figure 3A/C**). As expected, the genome of all the tested mutant cell lines contains two *Kpn*I bands-containing GK whose size is increased by ∼800 bp, corresponding to the length of the repair cassette. It is noteworthy that, following cleavage by Cas9 in the same locus, it is likely that several GK copies were deleted by homologous recombination. Indeed, a Southern blot analysis of genomic DNA digested by *Mfe*I (which is absent in the GK repeat units), revealed a significant reduction of the size of one GK allelic cluster in the 1B10, 2G2 and 2F11 clones (**Figure S3**). As expected, a western blot analysis with the anti-GK immune serum showed that GK expression is abolished in all the analyzed mutant cell lines, confirming that they are *bona fide* GK null mutants (GK^−/−^) (**Figure 3D**). In addition, glycerol metabolism is abolished in the GK^−/−^ cell lines, as shown by quantitative proton NMR spectrometry analyses of the ^13^C-enriched end products excreted from the metabolism of uniformly ^13^C-enriched glycerol ([U-^13^C]-glycerol) (Bringaud et al., 2015, Pineda et al., 2018, Allmann et al., 2021). Indeed, the parental PCF *T. brucei* convert [U-^13^C]-glucose or [U-^13^C]-glycerol to ^13^C-enriched acetate and succinate. In contrast, the excretion of ^13^C-enriched end products from [U-^13^C]-glycerol is abolished in the GK^−/−^ 2E6 cell line, while the metabolism of [U-^13^C]-glucose is unaffected (**Figure 3E**). Taken together, these data demonstrate the high efficiency of a single transfection with gRNA/*Sp*Cas9 complexes to inactivate all copies of a large multigene family.

**Figure 3-.**
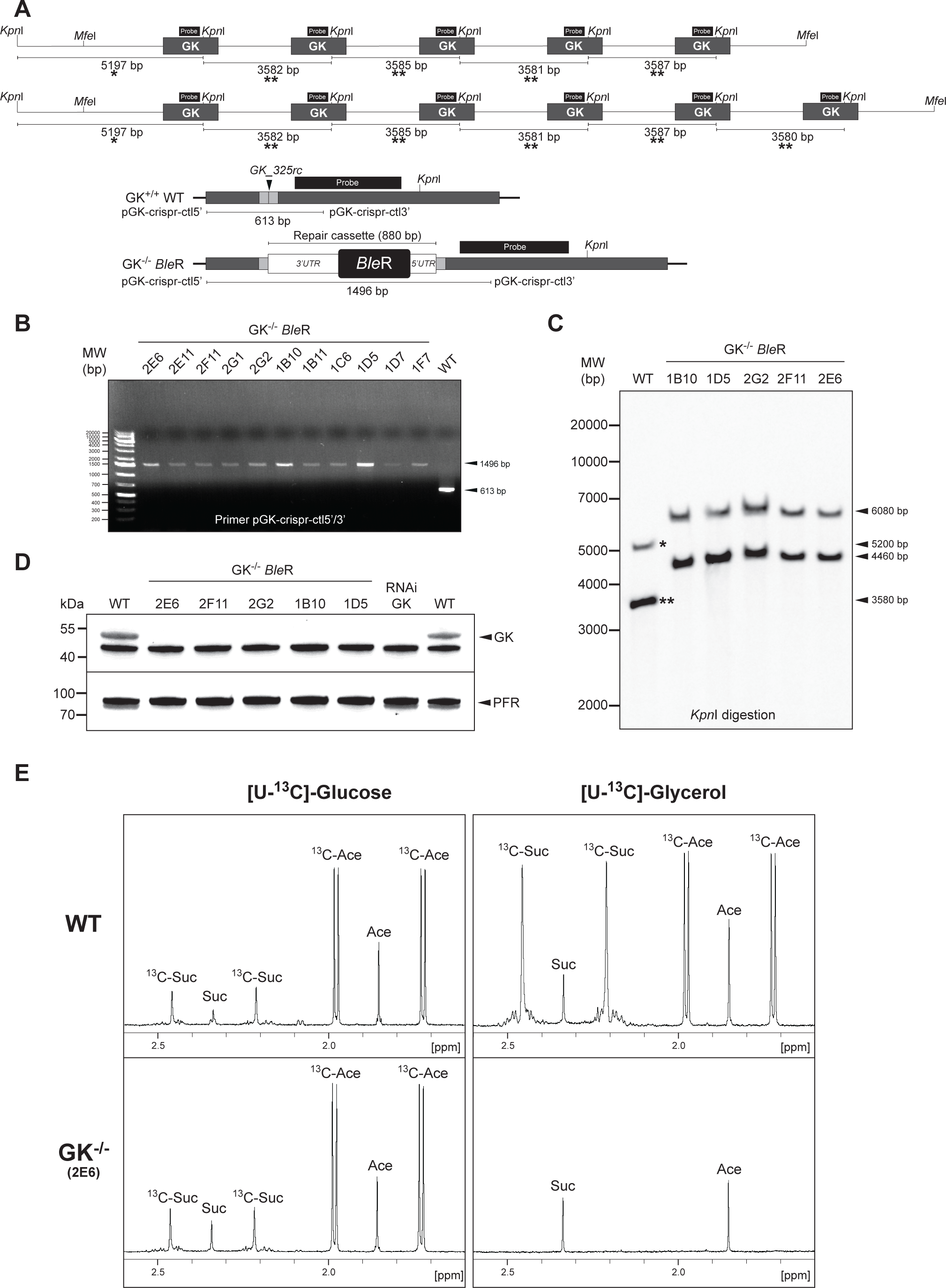
Inactivation of the multigenic family encoding the glycerol kinase (GK) in *T. brucei* PCF. **(A)** Schematic representation of the two alleles of the GK family in *T. brucei* and the inactivation strategy used by insertion of the phleomycin resistance marker *Ble*R. The position of the guide RNA is indicated by a vertical arrow (GK_325rc) and the 50-bp flanking sequences allowing repair through HDR are shown in gray. The position of the probe used for the Southern-blot analysis is indicated by a black box. **(B)** Confirmation by PCR of GK gene inactivation on both alleles in PCF. PCR products from various phleomycin-resistant clones (primer pGK-crispr-ctl5’/pGK-crispr-ctl3’) are analyzed on an agarose gel. **(C)** Southern-blot analysis of various phleomycin-resistant clones. The two bands detected in the parental cells (WT) correspond to the GK copy located at the 5’ extremity of the clusters (one asterisk) and to all the other GK copies (two asterisks). The insertion of the resistance marker increases the size of both of these bands by 880 bp in the phleomycin-resistant clones. **(D)** Western blot analysis of whole-cell extracts from different phleomycin-resistant *T. brucei* PCF clones. RNAi targeting GK was included as a control (Pineda et al., 2018). Antibodies against the paraflagellar rod (PFR) were used as a loading control. **(E)** ^1^H-NMR analysis of ^13^C-enriched end products (succinate and acetate, ^13^C-Suc and ^13^C-Ace, respectively) excreted from the metabolism of [U-^13^C]-glucose and [U-^13^C]-glycerol, by the parental (WT) and clone 2E6 (GK^−/−^) PCF cells. A portion of each spectrum ranging from 1.6 ppm to 2.6 ppm is presented.

### *Sp*Cas9 is also functional in other trypanosomatids

We also tested the efficiency of RNP complex delivery to edit the genome of the promastigote forms of *Leishmania infantum* and of the bloodstream and procyclic forms of *Trypanosoma congolense*. Several approaches have been used to inactivate the gene encoding ALDH, the mitochondrial enzyme responsible for converting acetaldehyde to acetate in *L. infantum* promastigotes (LINF_250017300, ALDH, manuscript in preparation), using an *Sp*Cas9/gRNA complex targeting the ALDH sequence and various repair cassettes. Here we used repair cassettes to insert (i) a puromycin resistance marker (*Pac*R), (ii) a fluorescent protein (monomeric RED, mRED) and (iii) a short sequence containing stop codons and a *Bam*HI restriction site (SBS) as described above (**Figure 4A**). After transfection, clones were selected either by addition of puromycin (*Pac*R cassette, ALDH^−/−^ PAC cells), by cell cytometry at 595/613 nm (mRED cassette, ALDH^−/−^ mRED cells) or by PCR and sequencing after cell cloning (SBS cassette, ALDH^−/−^ SBS cells). In each condition, the insertion of the repair cassette was checked by PCR with primers flanking the insertion site (**Figure 4A/B**). We were able to obtain homozygous mutant clones for each repair cassette, with efficiencies ranging from 30% (ALDH^−/−^ SBS cells) to 83% (ALDH^−/−^ mRED cells) (**Table 1**). It should be noted that the *Pac*R and mRED cassettes are only composed of the corresponding ORF inserted in frame with the ALDH coding sequence (**Figure 4A**). The mRED protein showed a mitochondrial-like pattern by immunofluorescence in the ALDH^−/−^ mRED cell line, as opposed to the cytosolic-like pattern observed for mRED expressed with an expression vector, which suggests that the N-terminal mitochondrial targeting motif of ALDH targeted the chimeric ALDH/mRED protein to the mitochondrion (**Figure 4C**). To confirm that both ALDH alleles were indeed inactivated, we quantified the product of the ALDH enzymatic reaction, *i.e.*, acetate, which is excreted in the medium from the metabolism of threonine. As expected, production of ^13^C-enriched acetate is abolished in the ALDH^−/−^ PAC clone 1B1, as shown by proton NMR spectrometry analysis of the ^13^C-enriched end products excreted from the metabolism of [U-^13^C]-threonine (**Figure 4D**) (Bringaud et al., 2015). Integrating an ectopic copy of the ALDH encoding gene in an ALDH^−/−^ PAC cell line restored ^13^C-enriched acetate production from the metabolism of [U-^13^C]-threonine, as opposed to expression of GFP (**Figure 4D**). Similarly, the ALDH^−/−^ SBS mutant (clone A1) no longer excretes ^13^C-enriched acetate from the metabolism of [U-^13^C]-threonine (**Figure 4D**).

**Figure 4-.**
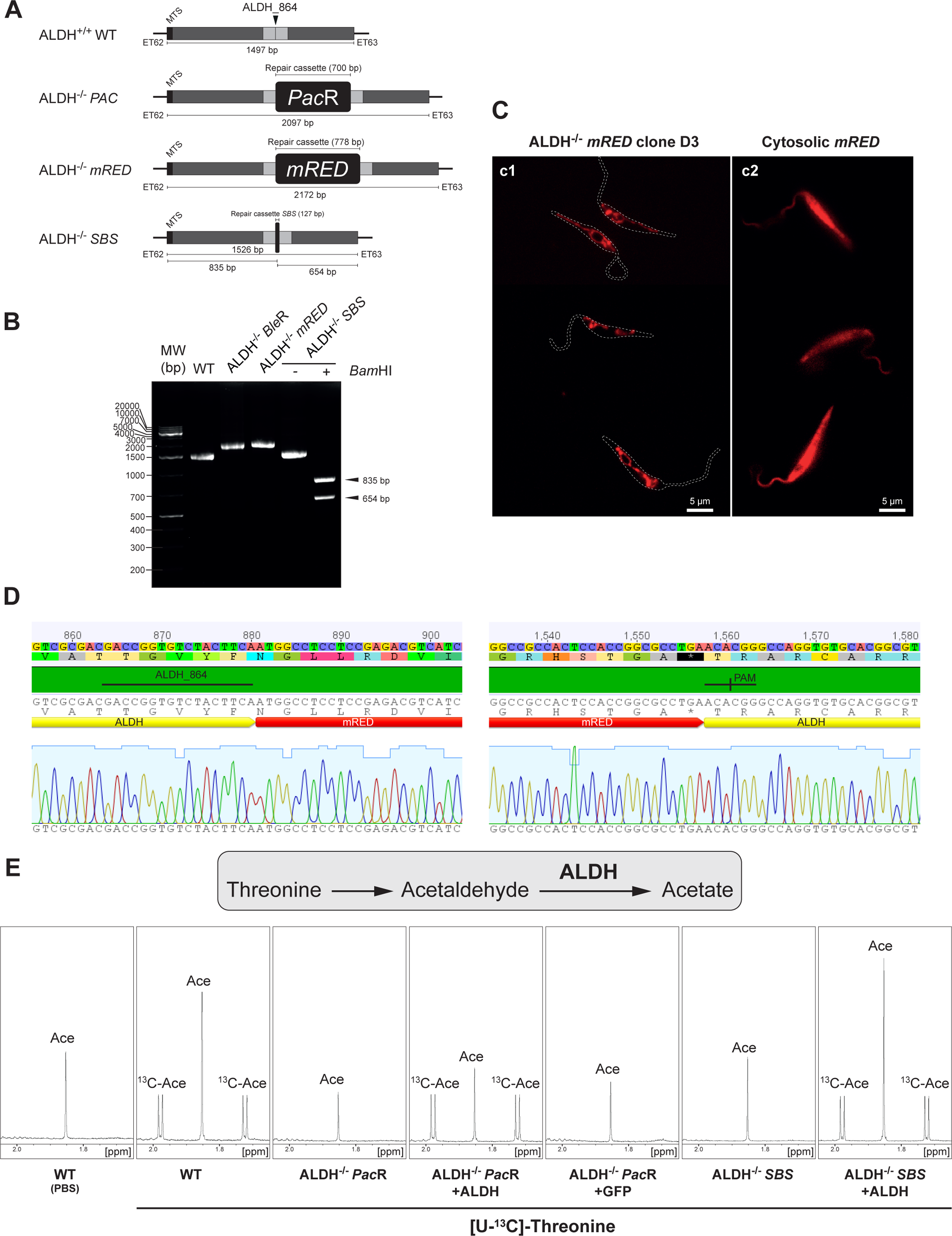
ALDH inactivation in *L. infantum* promastigote. **(A)** Schematic representation of the ALDH locus and the three inactivation strategies used, *i.e.*, insertion of the puromycin resistance marker (*Pac*R), of the monomeric RED fluorescent protein (*mRED*) or of a short sequence containing a succession of stop codons (SBS). The position of the guide RNA is indicated by a vertical arrow (ALDH_864) and the 50-bp flanking sequences allowing repair through HDR are shown in gray. **(B)** Confirmation by PCR of ALDH gene inactivation on both alleles. PCR products from phleomycin-resistant, RED fluorescent and SBS clones (primer ET62/ET63) are analyzed on an agarose gel. In the case of the SBS strategy, the PCR product was digested by *Bam*HI. **(C)** The chimeric ALDH/mRED protein is expressed in the mitochondrion. The subcellular localization of ALDH/mRED in the ALDH^−/−^ D3 clone (c1) was revealed by mRED fluorescence and compared to the cytosolic expression of mRED (c2). **(D)** Portion of the chromatogram showing the fusion of the mRED sequence with the ALDH sequence. **(E)** ^1^H-NMR analysis of ^13^C-enriched end products (^13^C-acetate, ^13^C-Ace) excreted from metabolism of [U-^13^C]-threonine metabolism by the parental (WT), ALDH^−/−^ *Pac*R clone 1B1 re-expressing (ALDH^−/−^ *Pac*R +ALDH) or not the ALDH genes and ALDH^−/−^ SBS clone A1 re-expressing (ALDH^−/−^ *Pac*R +ALDH) or not the ALDH gene. A rescue control was performed by expressing GFP in the ALDH^−/−^ *Pac*R clone. A portion of each spectrum ranging from 1.65 ppm to 2.05 ppm is presented.

To test this approach in *T. congolense*, we targeted the *LysoPLA* gene (TcIL3000.A.H_000623300), which is a non-essential gene encoding an excreted lysophospholipase in *T. brucei* (Monic et al., 2022, Tounkara et al., 2021). Repair cassettes containing either the phleomycin-resistant gene (*Ble*R) flanked by 5’ and 3’ regulatory sequences or the marker-free SBS sequence were used to transfect BSF and PCF, respectively (**Figure 5A**). As above, the insertion of the repair cassettes was checked by PCR and sequencing from genomic DNA isolated from Phleomycin selection (*Ble*R cassette) or cell cloning (SBS cassette). The four BSF clones tested are all homozygous mutants, further confirmed by western-blot analyses using an anti-*Lyso*PLA immune serum showing that *Lyso*PLA is no longer expressed (**Figure 5A/B/C**). Similarly, three out of the five PCF clones tested are homozygous mutants (**Figure 5D**, clones 1B6, 1E10 and 2C3), as confirmed by western-blot analyses (**Figure 5E**).

**Figure 5-.**
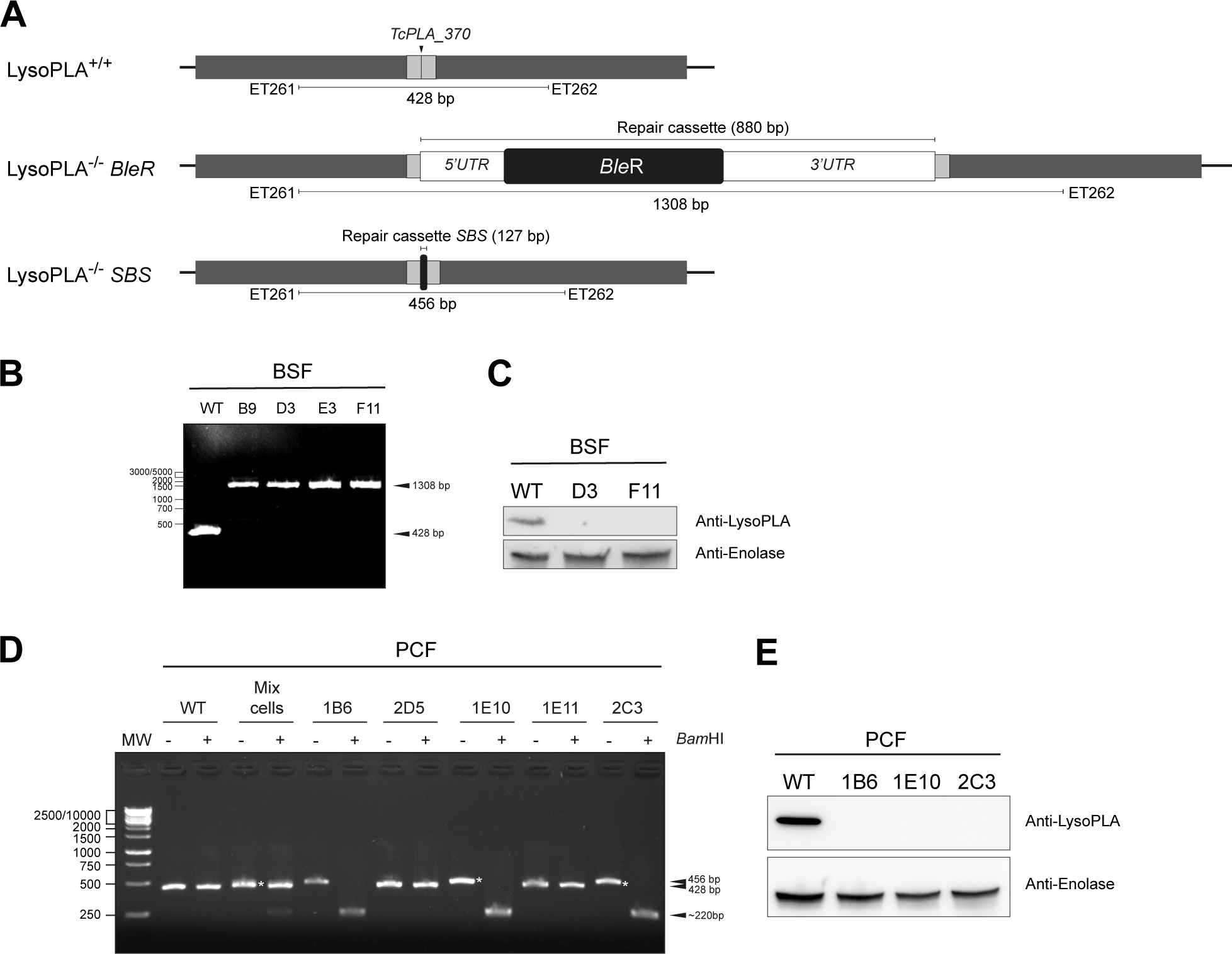
Inactivation of the *Lyso*PLA in *T. congolense* BSF and PCF. **(A)** Schematic representation of the *LysoPLA* locus in *T. congolense* and the two inactivation strategies used, *i.e.*, insertion of the phleomycin resistance marker (*BleR*) or a short sequence containing a succession of stop codons (SBS). The position of the guide RNA is indicated by a vertical arrow (*TcPLA_370*) and the 50-bp flanking sequences allowing repair through HDR are shown in gray. **(B)** Confirmation of *LysoPLA* gene inactivation on both alleles in *T. congolense* BSF cells, by PCR analysis of various phleomycin-resistant clones (primer ET261/ET262). **(C)** Western blot analysis of whole-cell extracts from two phleomycin-resistant *T. congolense* BSF clones. Antibodies against enolase were used as a loading control. **(D)** Confirmation of *LysoPLA* gene inactivation on both alleles in *T. congolense* PCF cells by agarose gel analysis of *Bam*HI-digested PCR products from various phleomycin-resistant clones (primer ET261/ET262). The “Mix cells” lane corresponds to the cell population before cloning, containing parental cells, and heterozygous and homozygous mutants. The asterisk indicates the 456-bp band, which does not contain a *Bam*HI restriction site. Clones 2D5 and 1E11 exhibit the wild-type profile. **(E)** Western blot analysis of whole-cell extracts from marker-free (SBS) *T. congolense* PCF clones. Antibodies against enolase were used as a loading control.

We also used *Sp*Cas9 to inactivate the GK multigene family in *T. congolense*, which is composed of 3 GK copies per allele, one of which is a pseudogene with two frameshifts (**Figure 6A**, TcIL3000_0_55170, TriTrypDB). BSF were transfected with a *Sp*Cas9/gRNA complex and a repair *Ble*R cassette containing 5’ and 3’ regulatory sequences. Among the four clones tested, three were homozygote mutants and one was heterozygote (**Figure 6B**). The absence of GK expression was confirmed by western blot analyses using anti-GK antibodies (**Figure 6C**) and by quantitative proton NMR spectrometry analyses of excreted end products from the metabolism of [U-^13^C]-glycerol and glucose (**Figure 6D**). Indeed, the production of ^13^C-enriched succinate and acetate from the metabolism of [U-^13^C]-glycerol is abolished in a mutant cell line, while the conversion of non-enriched glucose to succinate and acetate is not affected (**Figure 6D**). In conclusion, our data clearly showed that transfection of RNP complexes containing *Sp*Cas9 has the capacity to rapidly and efficiently modify all members of multigene families in *T. congolense* and probably all trypanosomatids.

**Figure 6-.**
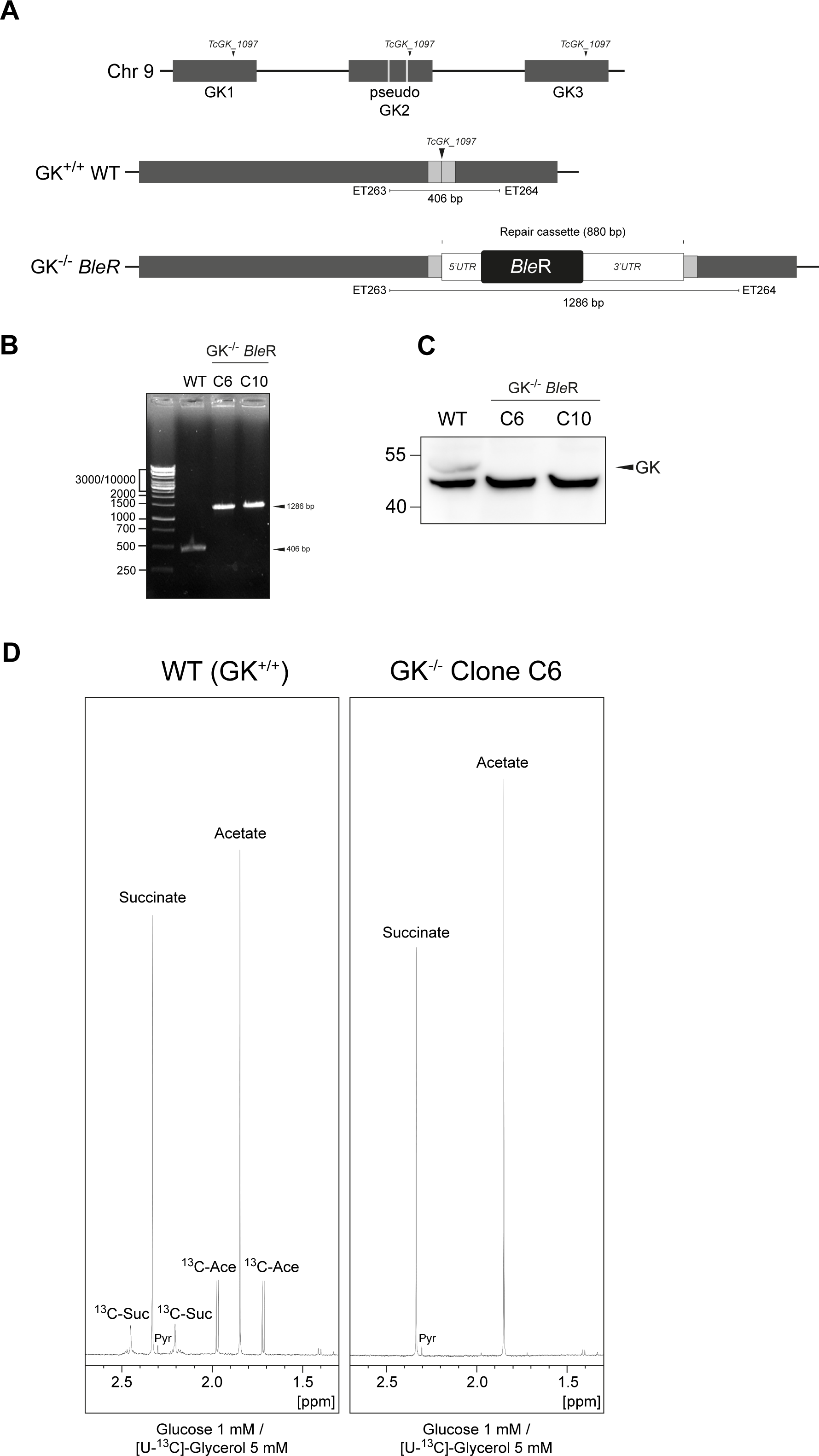
Inactivation of the multigenic family encoding the glycerol kinase (GK) in *T. congolense* BSF. **(A)** Schematic representation of the GK locus and its inactivation by inserting the phleomycin resistance marker *Ble*R. The position of the guide RNA is indicated by a vertical arrow (*TcGK_1097*) and the 50-bp flanking sequences allowing repair through HDR are shown in gray. **(B)** PCR confirmation of GK gene inactivation on both alleles in two different BSF clones. **(C)** Western blot analysis of whole-cell extracts from two phleomycin-resistant *T. congolense* BSF clones. **(D)** ^1^H-NMR analysis of end products (succinate and acetate) excreted from the metabolism of glucose and [U-^13^C]-glycerol by the parental (WT) and clone C6 (GK^−/−^) BSF cell lines. A portion of each spectrum ranging from 1.3 ppm to 2.7 ppm is presented. In the NMR experiments, *T. congolense* BSF were incubated in the presence of a mixture of D-glucose (1 mM) and D-[U-^13^C]-glycerol (5 mM) to keep them alive. Resonances were assigned as follows: Ace, acetate; ^13^C-Ace, ^13^C-enriched acetate; Suc, succinate; ^13^C-Suc, ^13^C-enriched succinate; Pyr, pyruvate.

## DISCUSSION

Since 2014, kinetoplastid studies using CRISPR/Cas9 technology have all employed parasites that constitutively express Cas9, which required genome integration of the RNP-encoding gene, even gRNAs and in some cases. However, these potentially compromise parasite growth (Peng et al., 2014). More recently, Soares Medeiros *et al*. demonstrated that transfection of the Cas9-gRNA RNP complex also induced rapid and efficient genome editing in kinetoplastids, but only with a Cas9 derived from *Staphylococcus aureus*, and hypothesized that *Sp*Cas9 was too large to be transfected and functional (Soares Medeiros et al., 2017). In this current study, we demonstrate that *Sp*Cas9 is fully functional and effective after transfection into the parasites. We have shown that both the commercial *Sp*Cas9 (IDT) and the *Sp*Cas9 produced in our laboratory enable rapid and efficient genome modification in several kinetoplastids (*T. brucei*, *T. congolense and Leishmania*), across different life cycle stages (insect and mammalian stages). We were able to target single genes and multigene families.

This approach based on transfection of RNP complexes offers several advantages including, (i) adaptability to any laboratory and field strains, (ii) for non-essential genes, both alleles are inactivated simultaneously, which is also valid for large multigene families, (iii) Cas9 remains transiently in the transfected cell, preventing them from the deleterious effect of constitutive Cas9 expression, and (iv) no need for selection markers, which implies that numerous modifications can be achieved in the same cell line. Peng *et al*. demonstrated that mutations induced by Cas9 were mediated by MMEJ, a process that results in a deletion between homologous regions (Peng et al., 2014). We have also observed such deletions in *T. brucei* with both the commercial *Sp*Cas9 (from IDT) and laboratory-produced *Sp*Cas9, in the absence of repair cassettes. This approach proved to be very efficient since we inactivated the GFP gene introduced in *T. brucei* PCF with an efficiency close to 50%, all within a few days. When double-strain breaks through Cas9 are combined with repair cassettes, it becomes easy to achieve targeted insertions through HDR, allowing inactivation or tagging of genes at their endogenous loci. We have thus been able to insert resistance markers (*Ble*R and *Pac*R), a gene encoding the RED fluorescent protein, or a short insertion sequence containing a series of stop codons enabling the inactivation of the target gene. We did not test the limit for the homology arms’ size, but in *T. cruzi* and *Leishmania*, approximately 30 bp is sufficient (Soares Medeiros et al., 2017). Finally, the insertion of short coding sequences at the 5’ and/or 3’ ends of the targeted genes (in order to tag them) can be achieved on both alleles without a selection marker and without significant modification of the UTRs (Morel et al., 2023). In terms of efficiency, there appears to be no set rule, i.e., this may depend on the gRNA, the targeted gene, the stage of division, and how the parasites are selected. However, adding regulatory sequences upstream and downstream of the insertion cassette encoding a resistance marker, appears to enhance the selection of homozygous clones for *T. brucei* BSF. We did not test the addition of regulatory sequences in *Leishmania* since the obtained homozygote rate was already very high (**Table 1**). In *T. cruzi*, successive transfections appear to significantly increase efficiency (Soares Medeiros et al., 2017), and this is an interesting approach when homozygous clones are not obtained. Here, we consistently obtained homozygous clones for non-essential genes. However, for genes suspected to be essential, only heterozygous clones are obtained. Therefore, successive transfections should enable the confirmation of their essentiality, if only heterozygous clones are obtained. It should be noted that successive transfections may also lead to the selection of chromosomal polysomy (Tovar et al., 1998).

Another important point to consider is the optimization of Cas9 for its importation by electroporation. Soares Medeiros *et al*. reported that recombinant *Sp*Cas9 is not active on *T. cruzi*, which was interpreted as non-internalization by electroporation due to a size issue (Soares Medeiros et al., 2017). In agreement with their hypothesis, they showed the functionality of a smaller Cas9 (Cas9 from *Staphylococcus aureus*), which is abolished by fusion with GFP (Peng et al., 2014). However, the recombinant *Sp*Cas9 is functional in the same experimental set up. The only difference between the recombinant *Sp*Cas9 used by Soares Medeiros *et al*. and us it the presence of 2 and 3 NLS sequences, respectively. These data suggest that the size of the ribonucleoprotein complex is not a limiting factor, however the number of NLS sequences to achieve effective nuclear targeting seems to be an important factor to consider. Very recently, Minet *et al*. also managed to transfect and modify *T. congolense* bloodstream forms using a commercial *Sp*Cas9, which contains 3 NLS in its sequence, further demonstrating that this protein is fully functional in kinetoplastids (Minet et al., 2023).

Finally, this system is likely functional in all cells that can be transfected, enabling more relevant studies on field strains. Indeed, this system is also valuable for studying cells that are difficult to cultivate (low cell density) and that previously required a significant number of cells for transfection through classical homologous recombination approaches. We used 5×10^5^ cells to efficiently edit both alleles of the targeted genes, but we believe that this approach could be adapted to many fewer cells, provided that a sufficient number of clonal cells are sorted by flow cytometry. The transfection of the RNP-gRNA complex and its various derivatives, including dead Cas9, dCas9-methyltransferases, activators, etc., presents a wealth of exciting research opportunities across diverse cell types (Engstler and Beneke, 2023, Gomaa et al., 2022). Notably, this innovative approach holds great promise for advancing our understanding of biological processes showing several redundant pathways, such as some metabolic capacities (Millerioux et al., 2018, Wargnies et al., 2018), for which the need to inactivate multiple enzymes has often been hampered by the scarcity of selection markers.

## EXPERIMENTAL PROCEDURES

### Trypanosomes and cell cultures

The procyclic forms (PCF) of *T. brucei* EATRO1125.T7T (TetR-HYG-T7RNAPOL-NEO, where TetR stands for tetracycline resistance, HYG for hygromycin, T7RNAPOL for RNA polymerase T7, and NEO for neomycin) was cultured at 27°C with 5% CO_2_ in SDM79 medium containing 10% (vol/vol) heat-inactivated fetal calf serum, 5 µg/mL hemin, 0.04 mg/mL streptomycin, 40 U/mL penicillin (SigmaP4333), 25 µg/mL hygromycin and 10 µg/mL neomycin. The bloodstream forms (BSF) of *T. brucei* 427 90-13 (TetR-HYG-T7RNAPOL-NEO) was cultured at 37°C with 5% CO_2_ in Iscove’s modified Dulbecco’s medium (IMDM) supplemented with 10% (vol/vol) heat-inactivated fetal calf serum, 0.2 mM β-mercaptoethanol, 36 mM NaHCO_3_, 1 mM hypoxanthine, 0.16 mM thymidine, 1 mM sodium pyruvate, 0.05 mM bathocuproine, 1.5 mM L-cysteine, 5 µg/mL hygromycin and 2.5 µg/mL neomycin. *L. infantum* 263 promastigote form was cultured at 27°C with 5% CO_2_ in SDM79 medium containing 10% (vol/vol) heat-inactivated fetal calf serum, 5 µg/mL hemin, 0.04 mg/mL streptomycin and 40 U/mL penicillin (SigmaP4333). The BSF of *T. congolense* IL3000 was cultured at 34°C with 5% CO_2_ in MEM medium (Sigma M0643) containing 20% (vol/vol) heat-inactivated goat serum (InvitroGen 16210072), 6 mg/mL HEPES, 2 mg/mL NaHCO_3_, 1 mg/mL glucose, 100 µg/mL sodium pyruvate, 10 µg/mL adenosine, 14 µg/mL hypoxanthine, 4 µg/mL thymidine, 14 µg/mL bathocuproine, 2 mM glutamine and 0.2 mM β-mercaptoethanol, pH 7.2 to 7.4. The PCF of *T. congolense* was cultured at 27°C with 5% CO_2_ in MEM medium (Sigma M0643) containing 20% (vol/vol) heat-inactivated fetal calf serum, 6 mg/mL HEPES, 2 mg/mL NaHCO_3_, 5 µg/mL hemin, 2 mM glutamine and 8 mM Proline, pH 7.3-7.4. Growth was monitored by daily cell counting with the cytometer Guava® Muse® or Guava® *easyCyte*™.

### CRISPR/Cas9 inactivation

Gene inactivation was achieved by inserting double-stranded DNA corresponding either to a resistance marker (phleomycin or puromycin), to a gene encoding a fluorescent protein (monomeric RED), or to a short sequence containing 6 successive stop codons in the 3 reading phases and a *Bam*HI restriction site. These double-stranded DNA fragments were also flanked by 50 bp homologous to the 5’ and 3’ sequences of the Cas9 cut site. The EATRO1125.T7T PCF or 427 90.13 BSF (5×10^5^ cells) were respectively transfected, using Amaxa nucleofectorII, with 1 µg of purified cassette (phleomycin or puromycin resistance marker, mRED or Stop*Bam*HIStop), 30 µg of Cas9 protein from IDT preloaded with a mixture of TracrRNA (0.4 µmol) and gRNA (0.4 µmol). Cells were transfected using program X-001 or U-033 for *T. congolense* and selected or not with phleomycin (for *T. brucei* PCF 5 µg/mL or BSF 2.5 µg/mL and for *T. congolense* PCF 2.5 µg/mL or BSF 5 µg/mL) or puromycin (*T. brucei* PCF 1 µg/mL). Cells were cloned using a cell sorter (TBM Core facility), and the selection of inactivated cells was performed by DNA extraction using the NucleoSpin Blood kit (Macherey-Nagel) followed by PCR amplification using primers flanking the Cas9 cleavage site, see supplemental **Table S2**. Guide RNA were designed using EuPaGDT (Peng and Tarleton, 2015), from http://tritrypdb.org. Primers and guide RNA used were synthesized by Integrated DNA Technologies (IDT) and listed in supplemental **Table S2**.

### Southern-blot

A total of 2.5 µg of genomic DNA from T. *brucei* (EATRO1125.T7T) were subjected to *Kpn*I oe *Mfe*I digestion, electrophoresed in 0.8% agarose gel, blotted onto Hybond N^+^ membrane (Amersham), and hybridized with labelled probe at 50°C in 6X SSPE (1X SSPE: 0.18 mM NaCl, 10 mM NaH_2_PO_4_, 1 mM ethylenediaminetetraacetate, pH 7.0), 0.1% SDS and washed at 50°C using 0.5X SSPE-0.1% SDS, before revelation. Probes were obtained by PCR using the primers pGK-S55 and pGK-S53 (**Table S1**) and labelled with the PCR DIG Probe Synthesis Kit (Roche) according to the manufacturer and revealed using the DIG Luminescent Detection Kit and DIG Easy Hyb (Roche).

### Western-blot

Total protein extracts (5×10^6^ cells) were separated by SDS-PAGE (10%) and immunoblotted on TransBlot Turbo Midi-size PVDF Membranes (Bio-Rad). Immunodetection was performed using the primary antibodies, diluted in PBS-Tween-Milk (0.05% Tween20, 5% skimmed milk powder), rabbit anti-GK (1:1,000), rabbit anti-LysoPLA (1:1,000) and mouse anti-enolase (1:100,000, gift from P.A.M. Michels, Edinburgh, UK). Revelation was performed using a second antibody coupled to the HRP (anti-rabbit or anti-mouse IgG conjugated to horseradish peroxidase, Bio-Rad, 1:5,000 dilution) and detected using the Clarity Western enhanced-chemiluminescence (ECL) substrate as describes by the manufacturer (Bio-Rad). Images were acquired and analyzed with the ImageQuant Las 4000 luminescent image analyzer.

### Mitochondria staining on living cells

Rhodamine-123 (30 µg/mL) was added to cell culture (5×10^6^ - 1×10^7^ cells per mL) for 15 min at room temperature, then cells were washed twice with PBS and spread on slides. Images were acquired with MetaMorph software on Zeiss Axioplan 2 microscope and processed with ImageJ.

### Immunofluorescence

Cells were washed twice with PBS, then fixed with 2% paraformaldehyde (PFA) for 10 min at room temperature and 0.1 mM glycine was added for 10 min to stop the reaction. The cells were spread on slides and permeabilized with 0.05% triton X-100. After incubation in PBS containing 4% bovine serum albumin (BSA) for 20 min, cells were incubated for 1 h with primary antibodies diluted in PBS-BSA 4%, washed 4 times with PBS and incubated for 45 min with secondary antibodies diluted in PBS-BSA 4% followed by three washes. Kinetoplasts and nuclei were then labelled with DAPI (10 µg/mL) for 5 min. Slides were washed three times with PBS and mounted with SlowFade Gold (Molecular probes). Images were acquired with MetaMorph software on Zeiss Imager Z1 or Axioplan 2 microscope and processed with ImageJ.

### Analysis of excreted end-products from the metabolism of carbon sources by proton 1H-NMR

2 to 4×10^7^ *T. brucei* PCF, *Leishmania* promastigote or *T. congolense* BSF cells were collected by centrifugation at 1,400 x g for 10 min, washed twice with phosphate-buffered saline supplemented with 2 g/L NaHCO_3_ (pH 7.4) and incubated in 1 mL (single point analysis) of PBS supplemented with 2 g/L NaHCO_3_ (pH 7.4). Cells were maintained for 6 h at 27°C in incubation buffer containing one ^13^C-enriched carbon source (1 mM, [U-^13^C]-Glucose or [U-^13^C]-Glycerol or [U-^13^C]-Threonine; U stands for “uniformly ^13^C-labelled”), except for *T. congolense* BSF, which were incubated for only 1h30 at 37°C. The integrity of the cells during the incubation was checked by microscopic observation. The supernatant (1 mL) was collected and 50 µL of maleate solution in Deuterated water (D_2_O; 10 mM) was added as an internal reference. ^1^H-NMR spectra were performed at 500.19 MHz on a Bruker Avance III 500 HD spectrometer equipped with a 5 mm cryoprobe Prodigy. Measurements were recorded at 25°C. Acquisition conditions were as follows: 90° flip angle, 5,000 Hz spectral width, 32 K memory size, and 9.3 sec total recycle time. Measurements were performed with 64 scans for a total time close to 10 min 30 sec.

### Cas9 cloning, expression and purification Cas9

The *eSp*Cas9(1.1) gene containing two nuclear localization signals (NLS) was obtained from Addgene (Plasmid #71814) and cloned into the pST32 vector, which contains two N-terminal and C-terminal His-tag (Gift from Fanny Boissier, INSERM U1212 CNRS 5320, University of Bordeaux), using the *Nco*I and *Eco*RI restriction sites. A third nuclear localization signals (SV40) was added by hybridization of two complementary primers (**Table S1**) containing the SV40 NLS and cloned in frame at the 3’ end of the pST32-*eSp*Cas9(1.1) vector using *Eco*RI and *Xho*I restriction sites, generating the vector pST32-*eSp*Cas9(1.1)-3NLS (**Figure 1B**). The plasmid was then transformed into *E. coli* Rosetta 2(DE3) competent cells (Novagen).

A bacterial preculture was grown overnight at 37°C with shaking and used to inoculate 100 mL of LB-Miller medium (peptone 10 g, yeast-extract 5 g, NaCl 10 g, pH 7). The culture was grown at 37°C with shaking to an optical density at 600 nm (OD600) of 0.6 to 0.8. Protein expression was induced by adding isopropyl-ß-D-thiogalactopyranoside (IPTG; 100 µM), and the culture was kept at 18°C with shaking overnight. Cells were harvested by centrifugation, and the pellet was resuspended in 10 mL lysis buffer containing 500 mM KCl, 20 mM Hepes, 5 mM imidazole, pH 7.5 and protease inhibitor cocktail without EDTA (Merck). After lysis by sonication (20 sec, 4 times), the soluble fraction was obtained by centrifugation (30,000 x g, 30 min at 4°C) and purified by immobilized metal ion affinity chromatography (IMAC) using a His-Select Nickel Affinity Gel (Sigma) in a fast protein liquid chromatography (FPLC) system (ÄKTA; GE Healthcare Life Sciences). All chromatographic steps were performed at 4°C. Two mL of His-Select Nickel resin were equilibrated in buffer and packed in a XK 10/50 mm column housing (Omnifit) with 20 mL lysis buffer. The cleared lysate was loaded on the column using a syringe at 1 mL/min rate. The column with bound protein was washed first with buffer (20 mM Hepes, 500 mM KCl, 50 mM imidazole, pH 7.5) until the absorbance returned to baseline again. The protein was eluted by applying a gradient from 0% to 100% elution buffer (20 mM Hepes, 500 mM KCl, 1 M imidazole, pH 7.5) over 20 mL and collected in 2 mL fractions. All peak fractions were analyzed for the presence of *eSp*Cas9(1.1)-3NLS using SDS-PAGE, and the purity was estimated to be ∼80% based on band intensity (**Figure S1**). An alternative to the ÄKTA purification was to purified purify the Cas9 protein in batches using Ni^2+^-resin and incubated for 30 min at 4°C. The resin was washed three times with 50 mM imidazole in 500 mM NaCl, Tris-HCl pH 8 and 3 times with 250 mM imidazole in 500 mM NaCl, Tris-HCl pH 8. *eSp*Cas9(1.1)-3NLS was eluted with 500 mM imidazole in 500 mM NaCl, Tris-HCl pH 8. Fractions were analyzed by SDS-PAGE. The elution buffer was then exchanged for storage buffer (20 mM HEPES–KOH, 500 mM KCl, 1 mM DTT, pH 7.5) while concentrating the protein to a volume <1.5 mL using a 50,000 MWCO concentrator (Amicon) at 4,000 x g. Buffer exchange prevented precipitation in the concentrator. The concentrated fraction was then centrifuged for 10 min at 16,900 x g at 4°C to remove all precipitated material. The protein concentration was determined using a bicinchoninic acid (BCA) protein assay kit (Thermo Scientific), and the yield was determined to be approximately 5 mg/100 mL of bacteria culture.

### Statistical analysis

Experiments were performed at least in triplicates. Statistical analyses were performed using Prism (GraphPad) software. The results are presented as mean ± S.D. Where indicated the results were subjected to a two-sided student’s t-test to determine statistical differences against the indicated group (Confidence interval 95% - P-value style: 0.1234(ns); 0.0332 (*); 0.0021 (**); 0.0002 (***); <0.0001 (****)).

## ACKNOWLEDGMENTS

Cell sorter analyses were performed at the TBMCore facility (FACSility) on BD FACSAria™ III Sorter and we thank Atika Zouine and Vincent Pitard for technical assistance, data acquisition and interpretation (TBMCore CNRS 3427, INSERM US005, Université de Bordeaux). We also thank the CRISP’edit platform for their valuable advice during the setup of the CRISPR system. We thank Keith Gull (University of Manchester) for providing us the anti-PFR antibody. The Bringaud and Robinson teams are supported by the Centre National de la Recherche Scientifique (CNRS, https://www.cnrs.fr/), the Université de Bordeaux (https://www.u-bordeaux.fr/) and the Agence Nationale de la Recherche (ANR, https://anr.fr/) through the ParaFrap “Laboratoire d’Excellence” (LabEx, https://www.enseignementsup-recherche.gouv.fr/cid51355/laboratoires-d-excellence.html) (ANR-11-LABX-0024). The Bringaud team is also supported by the “Fondation pour la Recherche Médicale” (FRM, https://www.frm.org/) (“Equipe FRM”, grant n°EQU201903007845) and the ANR grant ADIPOTRYP (ANR19-CE15-0004-01) and the Robinson team by the ANR grant Structu-Ring (ANR-20-CE91-0003). M.O is the holder of a Canada research Chair and the recipient of a CIHR Foundation Grant. He was the holder of a University de Bordeaux IDEX fellowship.

## AUTHOR CONTRIBUTIONS

AC, HP, MP, OQ, MCA, BM, MS performed experiments and contributed to analysis and interpretation of the data. PMV, BM, RDR, OM, RL, BF conceptualized the study, contributed to analysis and interpretation of the data, TE performed experiments, conceptualized the study, contributed to analysis and interpretation of the data and wrote the manuscript, with also an input from all the authors.

## SUPPLEMENTAL MATERIAL

**Figure S1-.**
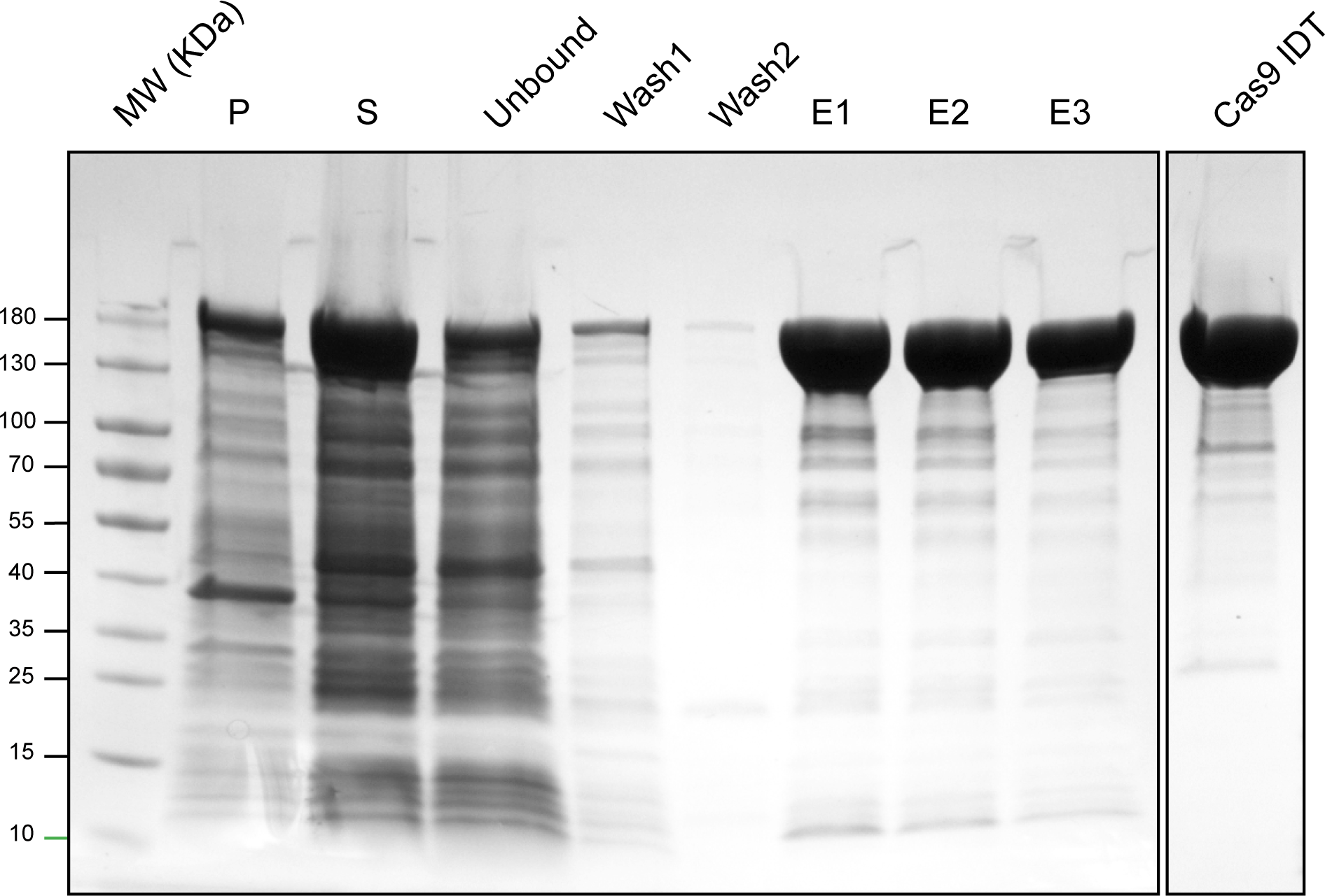
Expression and purification of the e*Sp*Cas9 from *E. coli*. The e*Sp*Cas9 protein expressed in *E. coli* was purified on a His-Select Nickel column. P, pellet; S, supernatant; Unbound, protein not retained on the column; Wash1 and 2 correspond to the wash fractions; E1 to E3 correspond to the elution fractions. The last lane represents 5 µg of purified Cas9 commercially available from IDT.

**Figure S2-.**
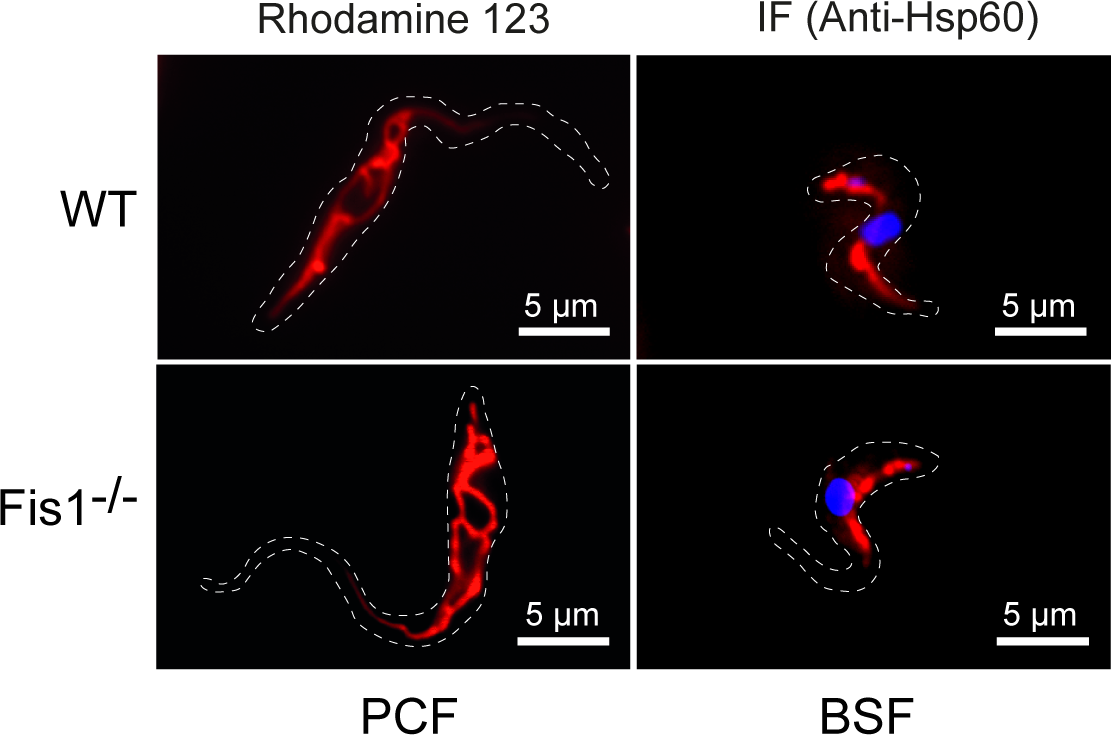
Inactivation of the *TbFis1* gene in both *T. brucei* PCF and BSF. Mitochondrial structure analysis using rhodamine 123 staining on living parental (WT) and *Tb*Fis1^−/−^ PCF cell lines, and with immunofluorescence using an antibody directed against Hsp60 to label and visualize mitochondrial shape in BSF cells. Only the BSF were labeled with DAPI.

**Figure S3-.**
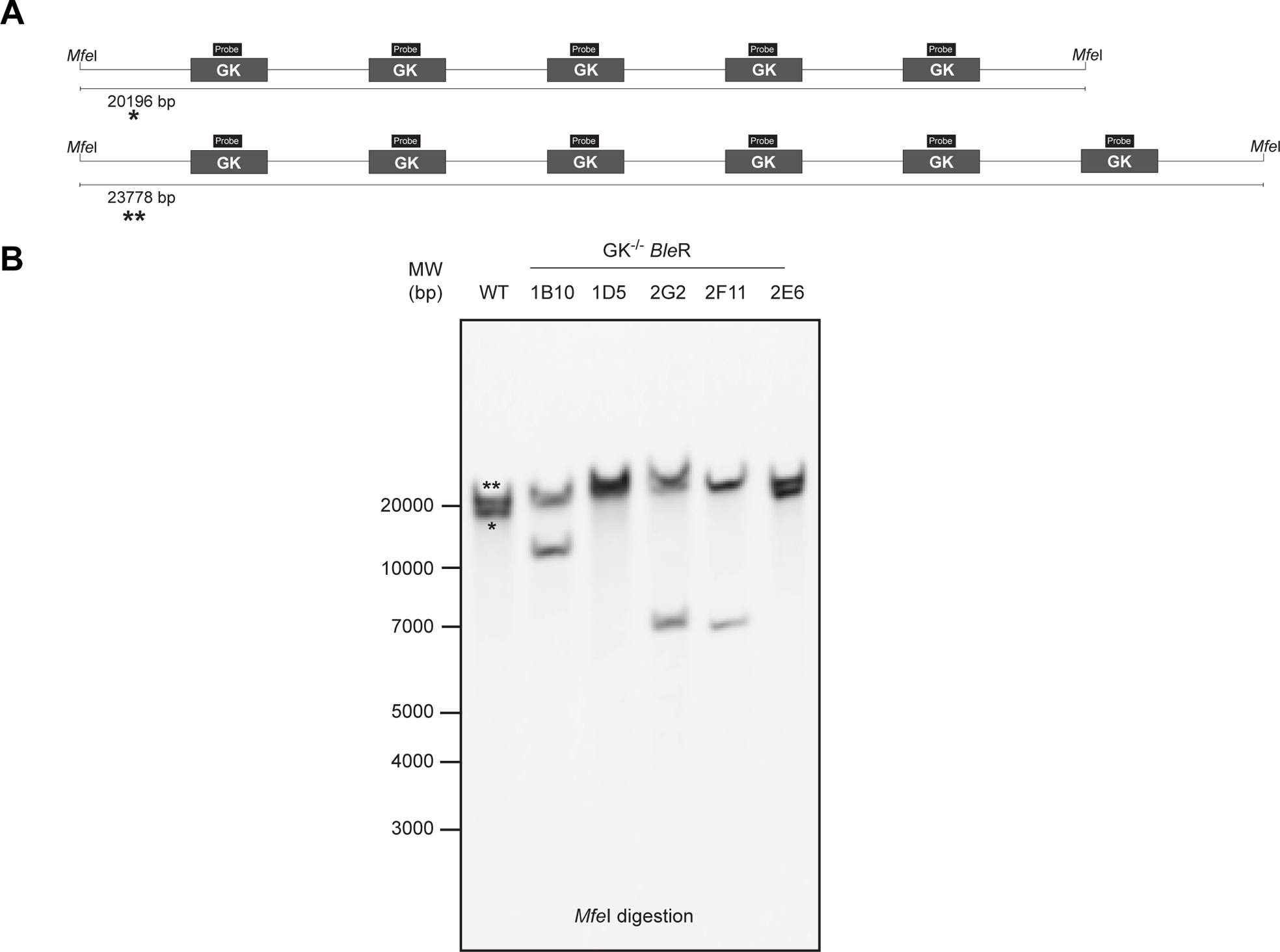
Inactivation of the multigenic family encoding the glycerol kinase (GK) in *T. brucei* PCF. **(A)** Schematic representation of the two alleles of the GK gene cluster in *T. brucei*. The position of the probe used for the Southern-blot analysis is indicated by a black box. **(B)** Southern blot analysis of various phleomycin-resistant clones. Genomic DNA was digested with *Mfe*I, separated on a 0.8% agarose gel, transferred to a membrane (Hybond-N), and probed with a PCR product labeled with the PCR DIG Probe Synthesis Kit (Roche). The two bands detected in the parental cell line (WT) correspond to the allele containing 5 (one asterisk) or 6 (two asterisk) copies of the GK gene. The insertion of the resistance marker into the various copies of GK can either increase the size of the *Mfe*I fragment or reduce it if deletion of GK copies occurred by homologous recombination after Cas9 cleavage.

**Table S1-.**
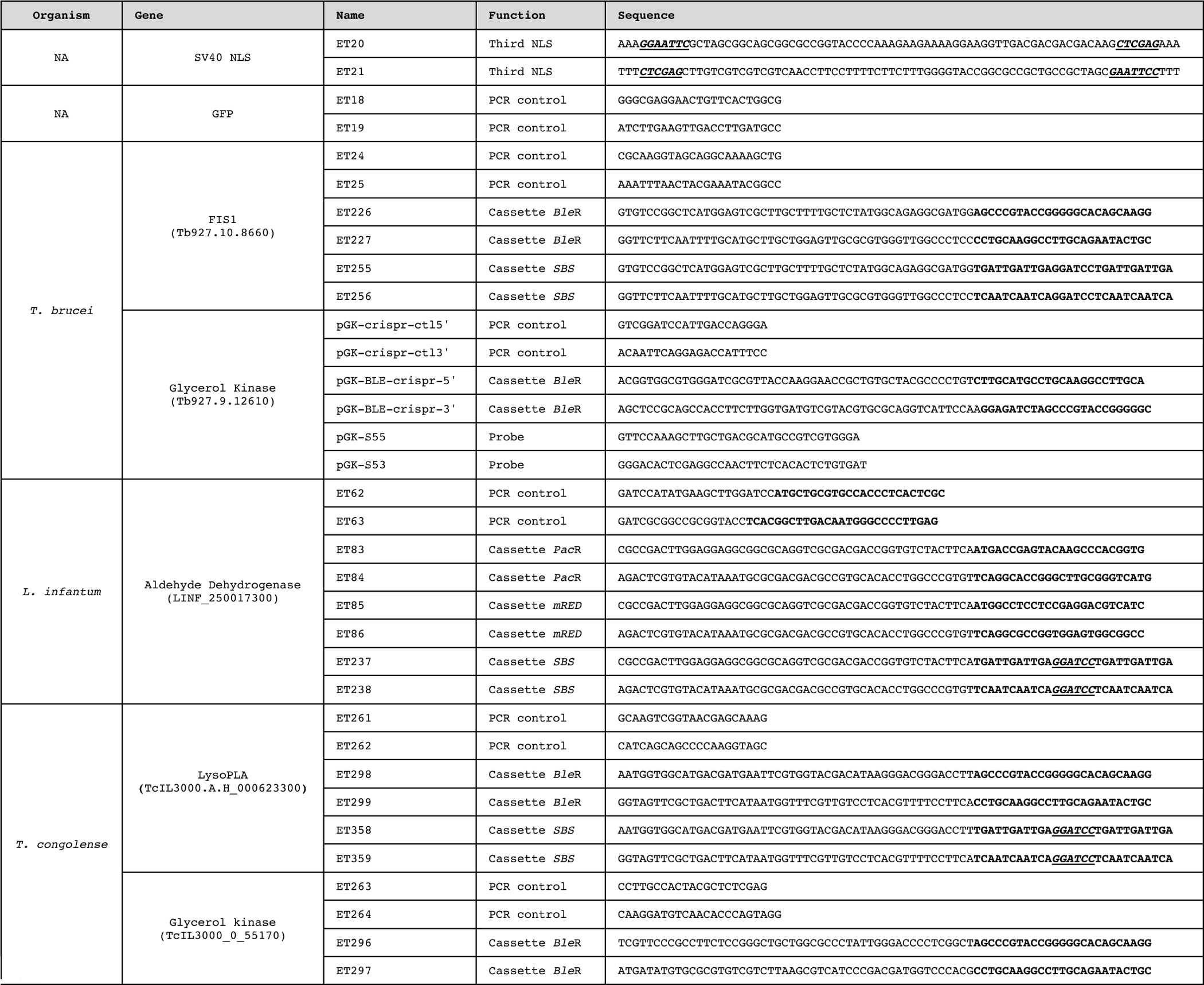
Oligonucleotides used.

**Table S2-.**
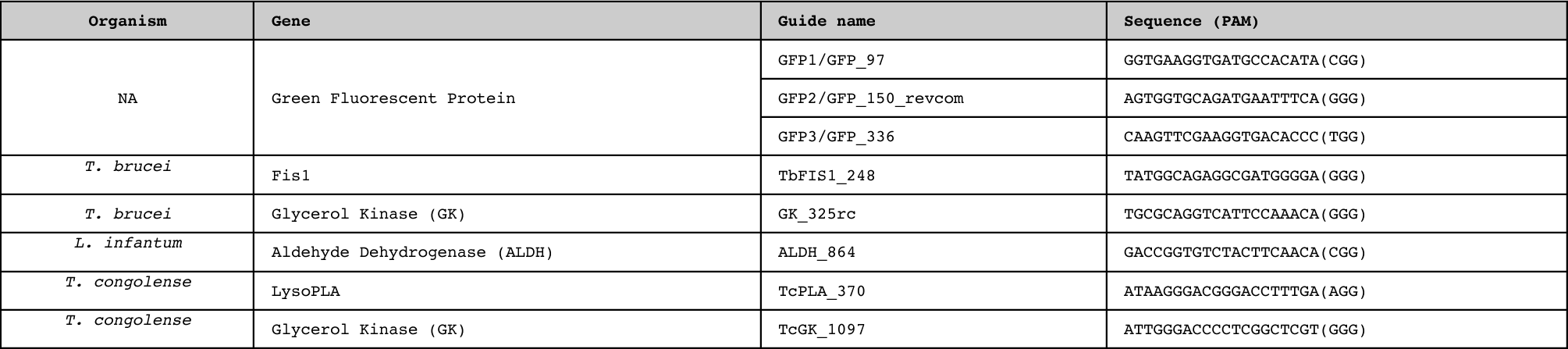
Guides RNA used.

